# dsSurvival 2.0: Privacy enhancing survival curves for survival models in the federated DataSHIELD analysis system

**DOI:** 10.1101/2023.05.25.542152

**Authors:** Soumya Banerjee, Tom R.P. Bishop

## Abstract

**Objective:** Survival models are used extensively in biomedical sciences, where they allow the investigation of the effect of exposures on health outcomes. It is desirable to use diverse data sets in survival analyses, because this offers increased statistical power and generalisability of results. However, there are often challenges with bringing data together in one location or following an analysis plan and sharing results.

DataSHIELD is an analysis platform that helps users to overcome these ethical, governance and process difficulties. It allows users to analyse data remotely, using functions that are built to restrict access to the detailed data items (federated analysis).

Previous works have provided survival modelling functionality in DataSHIELD (dsSurvival package), but there is a requirement to provide functions that offer privacy enhancing survival curves that retain useful information.

**Results:** We introduce an enhanced version of the dsSurvival package which offers privacy enhancing survival curves for DataSHIELD. Different methods for enhancing privacy were evaluated for their effectiveness in enhancing privacy while maintaining utility. We demonstrated how our selected method could enhance privacy in different scenarios using real survival data. The details of how DataSHIELD can be used to generate survival curves can be found in the associated tutorial.

## Introduction

Survival models are an important component of data science. They are used extensively in biomedical sciences to investigate the effect of exposures on health outcomes. Using data from diverse data sets is beneficial for increasing statistical power and generalisability of results. However, individual-level biomedical data are sensitive and ethical and legal considerations can make it challenging to move all the data required to one location or give analysts complete access to the data.

DataSHIELD is a platform that enables the non-disclosive analysis of distributed sensitive data. It permits analysts to work on remote datasets using functions that are designed to protect privacy through built in safeguards. Therefore the analysis can be run without sharing or fully accessing sensitive individual level data, which stay protected at their host institution [1].

We have previously developed a software package for DataSHIELD called *dsSurvival* 1.0 [2], which allows users to build survival models for each participating data set and meta-analyze hazard ratios while aiming to enhance the privacy of the data. In this work we have developed a new version of the package (*dsSurvival* 2.0) that adds significant new functionality to its previous version. We now allow the building and visualization survival curves for each data set that are designed to protect privacy.

## Main text

### Basics of survival models and survival curves

An important concept in survival models is a survival function:

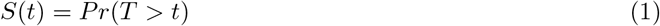

where *S*(*t*) is the survival function, *t* is the current time, and *T* is a random variable denoting time of death. *Pr()* is the probability that the time of death is greater than time *t* i.e. the probability of surviving until time *t*.

If we have observed survival times for a set of individuals, we can calculate the proportion surviving at each point during the study. These proportions can be plotted as a step function against time where the survival probabilities are constant between deaths. This is known as a survival curve and is a useful visualisation of survival data. For example, in biomedical research it might allow comparison of groups that had undergone different treatments.

### Architecture of DataSHIELD and dsSurvival

The DataSHIELD framework and *dsSurvival* have a client-server architecture. The communication between the client and a single server for *dsSurvival* 2.0 is shown in Fig. 1. A client-side general analysis function calls a server-side function. This function is executed on all the servers which store the data and summary results (that are designed to enhance privacy) are returned to the client. The server-side function has checks that ensure that privacy is enhanced in the results that are returned. The client-side function aggregates results from all servers and finally returns it to the analyst.

**Figure 1.**
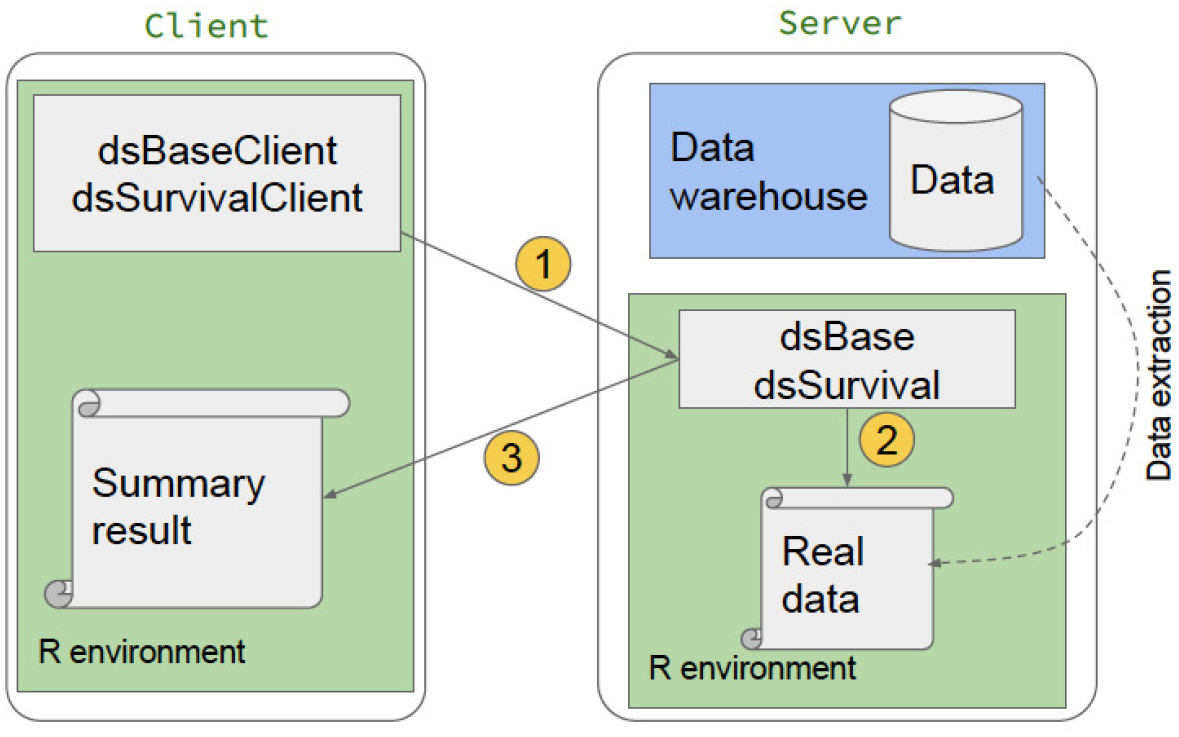
Example of workflow between a client and single data site in *dsSurvival*. Note that this can occur between a client and many sites. 1. User makes an analysis request in the DataSHIELD framework (e.g. generate a privacy enhancing survival plot) 2. The DataSHIELD function on the server side validates the request, and generates a result that is designed to enhance privacy (e.g. modified survival plot) 3. Result is returned to the client.

### Privacy and survival curves

A survival curve can be disclosive by revealing the time during which an event occurs. Each step in the curve corresponds to events for a group of individuals. If the size of the group is small (e.g. 1) then it becomes more likely that an adversary can attribute an event to an individual.

For example, if an adversary had a previously released survival curve, and an individual joined a study, a new drop in the curve could be attributed to that individual. The adversary would then know that the individual underwent an event, when it occurred and which particular subgroup the individual belongs to.

The objective of this work is to provide survival curves that aim to protect individuals privacy while still preserving useful information to the analyst. This provides novel functionality for the DataSHIELD platform.

### Methods for enhancing privacy in survival curves

Previous graphical approaches in DataSHIELD have used both deterministic anonymization and probabilistic anonymization. We investigated whether these techniques would be suitable for survival curves. Currently *dsSurvival* is designed to fit survival models per site, allowing users to meta-analyse these results to give an overall result. Similarly, our approach provides a survival curve per site, rather than a global survival curve.

Probabilistic anonymisation adds random noise to the data points before they are visualized. For survival curves noise added to the X-axis adds uncertainty to the time when the event occurred, and when added to the Y-axis obscures how many individuals undergo an event at a particular time. The amount of noise is specified as a percentage of the value it is being added to. We also ensure that even after noise is added, time (plotted on the X-axis in a survival curve) increases monotonically and proportion surviving (Y-axis in a survival curve) decreases monotonically. An example survival curve that has undergone probabilistic anonymisation is shown in Supplementary Section.

In deterministic anonymisation, the *k* nearest neighbours algorithm [3] is used to find centroids of the data points. Once the nearest neighbours are computed, the original data points are then moved to their centroids. This modified data is then plotted.

A review of existing literature suggests that two other options are available for generating privacy enhancing survival curves. The first of these is based around a smoothing technique (LOESS [Locally Estimated Scatterplot Smoothing]) [4]. Smoothing the steps in the survival curve makes it harder for an adversary to identify the timing of events and the number of individuals experiencing an event at a particular time. The smoothing process is achieved by fitting a low-degree polynomial to a subset of the data at each point. A smoothing parameter determines how much of the data is used to fit each polynomial - smaller values result in curves that track variations in the data closely and larger values give more smoothing.

The second method is to implement differentially private survival curves [5], which involves adding calibrated noise to the counts of events. While this makes strong promises about the protection of privacy, it is complex to implement. In particular it requires the calling framework to manage a privacy budget on behalf of each user which is depleted with each request, as the user learns more about the data. This is required for the promises around privacy protection to be upheld but this mechanism is not currently available in DataSHIELD.

We considered the different approaches available. Probabilistic anonymisation has the challenge that it is difficult to set the noise percentage to an appropriate amount to balance utility and privacy. Deterministic anonymisation has additional complexities when applied to skewed data such as survival data (more events happen at earlier times). The skewed data means that the nearest-neighbour algorithm “pulls” the data towards the centre of mass of the data.

While differential privacy is gaining traction in many applications, where a privacy budget can be managed, the lack of a solution to manage differential privacy in DataSHIELD meant it had to be ruled out.

We therefore choose to base our approach on LOESS, which has already been used in practice for survival curves [4].

Choosing a value for the smoothing parameter that would provide a suitable balance between privacy and utility could be challenging for data custodians. For example, a small parameter provides a curve closer to the true curve but risks compromising privacy. A larger parameter may smooth the curve too much such that it is less useful for research. Therefore we implemented LOESS using the the loess.as() function in the *fANCOVA* package [6], which automates the selection of the smoothing parameter. This is achieved by minimising a criterion that incorporates the number of variables in the model and the error in the fit of the model. We selected the corrected Akaike information criterion [AIC] because this offers better performance on smaller datasets [7].

### Ablation studies and sensitivity analysis

We assessed privacy risk in a publicly available dataset: the veteran dataset in the survival R package [8]. We show our privacy enhancing survival curves (generated using the automated LOESS smoothing procedure) on the full dataset (Fig. 2(a)) and the dataset with one patient removed (Fig. 2(b)). To demonstrate the effectiveness of the smoothing, we consider the individual that undergoes an event at time 61. In the unsmoothed curve this event is clearly visible and would identify that individual. In the smoothed curve, it is not possible to determine if an event occurred at time 61. While there is an additional inflection point at time *∼* 53 with the patient added, this does not match their actual event time of 61. This suggests it is very difficult to infer the characteristics of this single patient.

**Figure 2.**
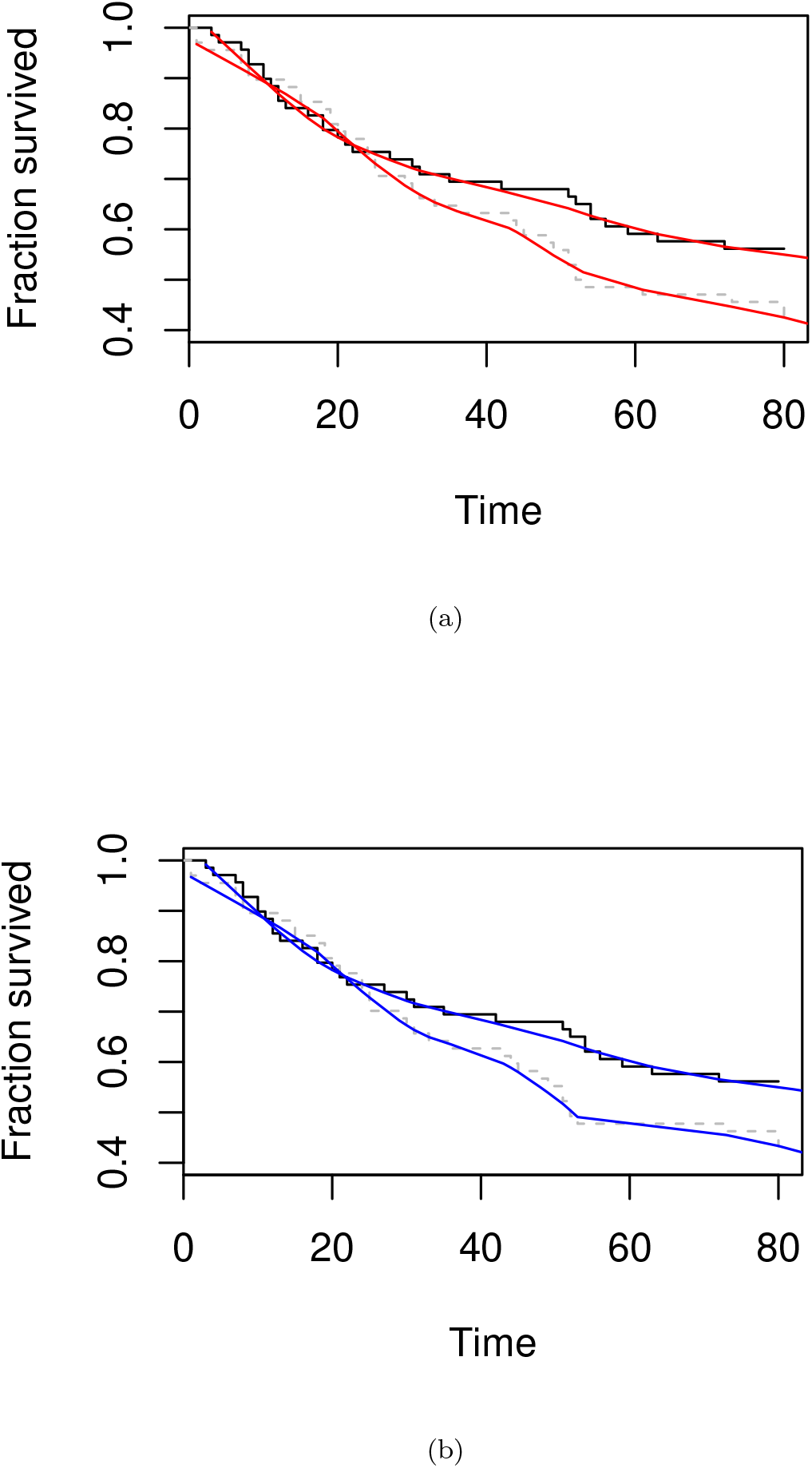
(a) Survival curve with LOESS smoothing using the *fANCOVA* package with all patients in the veteran dataset. The original unmodified survival curve is shown in black.(b) As per (a) but with one patient removed at time 61.

We also conducted ablation studies using this automated smoothing technique on a dataset where we randomly reduced the number of patients from 137 to 50 (ablation study). The resulting privacy enhancing survival curve is shown in Fig. 3(a). The survival curve with one additional patient removed is shown in Fig. 3(b). Additional survival curves generated from further ablation studies are available in the Supplementary Section.

**Figure 3.**
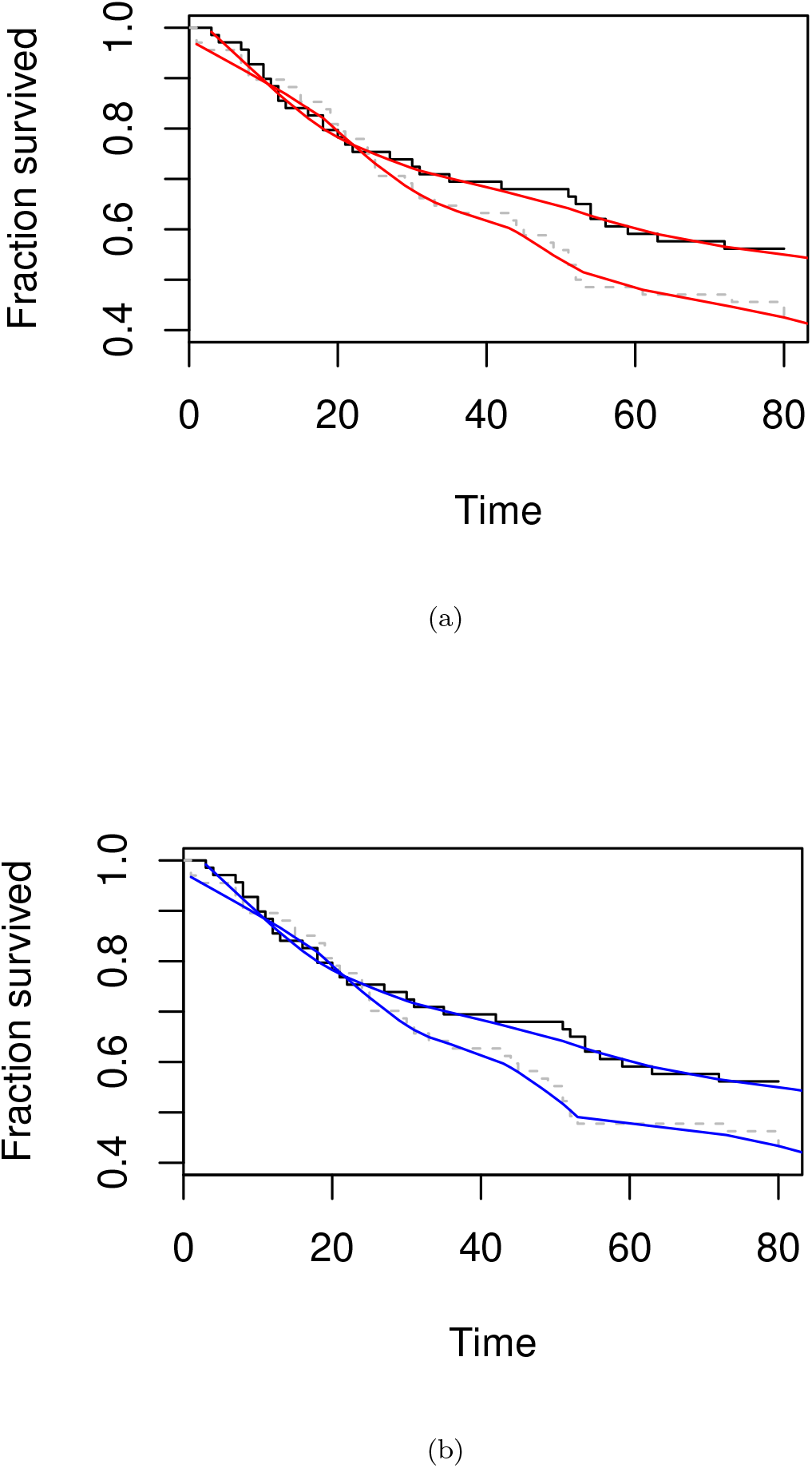
(a) Survival curve with LOESS smoothing using the *fANCOVA* package with randomly reduced number of patients (50 as opposed to 137) in the veteran dataset. The original unmodified survival curve is shown in black. (b) As per (a) but with one patient removed at time 61.

We conducted similar ablation studies to reduce the number of patients in synthetic data and observe the effect on survival curves (see Supplementary Section).

Based on our simulations and the fact that this technique was used in previous studies [4], we choose this automated smoothing technique as the primary implementation on *dsSurvival* 2.0 for generating survival curves.

### Summary of steps taken to enhance privacy

We summarise the steps we take to enhance privacy, noting that no procedure can completely remove disclosure risk. The smoothing procedure for survival curves ensures that it is very difficult to infer the precise time that an event takes place (for any given patient).

We modify the fraction of patients that survive (surv field) using LOESS procedure described above. In survival curves symbols show when the censoring events occur. These symbols are removed from the plot. This ensures that it is very difficult to infer the precise time that an event takes place (for any given patient).

Finally, we note that the DataSHIELD architecture also helps to minimize disclosure risk and protect privacy. The functions in DataSHIELD are designed to return summary aggregated statistics, enforcing requirements to enhance disclosure control.

### Computational pipeline and use case

We outline the development and code for implementing survival models and plotting of survival curves.

All code is available here in bookdown format with synthetic data: https://neelsoumya.github.io/dsSurvivalbookdown/

The computational steps are outlined below using synthetic data.

The initial steps outlined below create a Surv() object (available in the *survival* package) to generate the times and proportion surviving, which are then used as inputs to the survival curve.

#### Algorithm 1

Workflow for performing privacy enhancing survival analysis and plotting of survival curves.

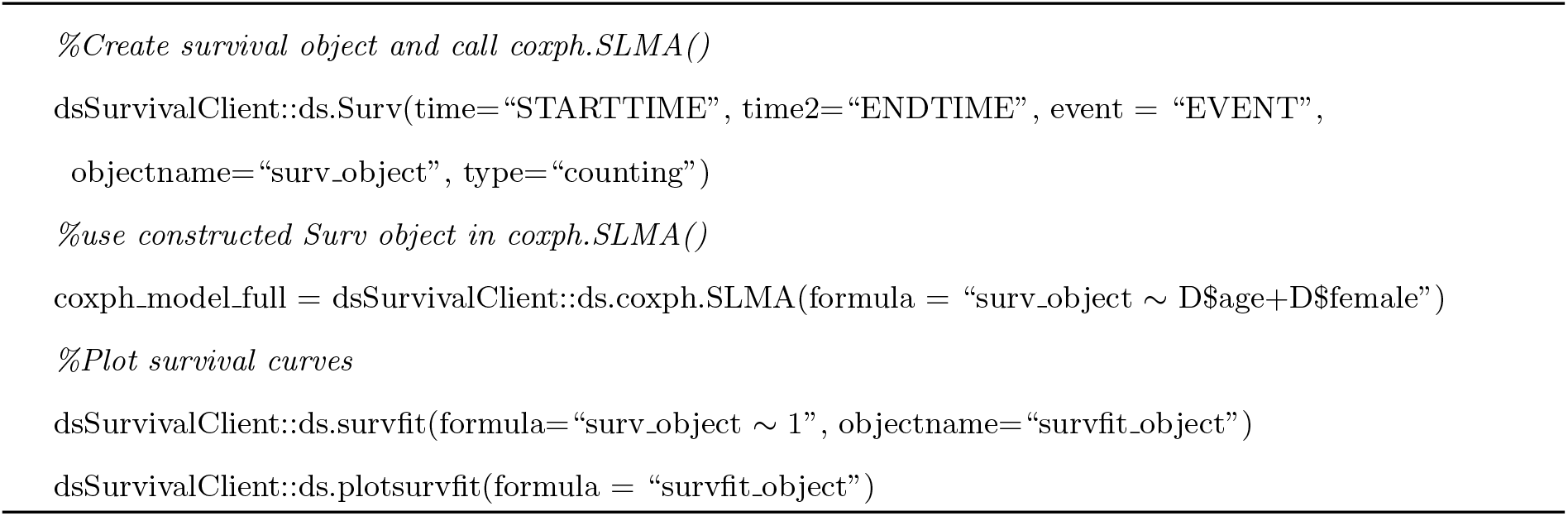

## Discussion

Our work adds to the existing *dsSurvival* package (which enables privacy enhancing meta-analysis of survival models in the DataSHIELD federated environment) by adding functionality to plot privacy enhancing survival curves. This will allow users to visually assess the differences in survival rates between groups, for example, while maintaining privacy.

We choose to build on the existing smoothing techniques [4] as this offered a pragmatic approach which we have adapted to a federated setting. Other methods such as probabilistic anonymisation are challenging to optimise the balance between privacy and utility.

Differential privacy has been used to ensure survival curves are privacy enhancing [5] and this is a promising area of future work [9]. Apart from the additional complexity in implementing the solution itself, there remain challenges around managing the privacy budget. This could be the subject of future work.

## Limitations

A limitation is that *dsSurvival* 2.0 provides a curve per study/dataset, and not a global curve. This might be possible with DataSHIELD in the future and would require secure exchange of values so that they can be ordered by time to build the survival curve. We also did not determine a minimum number of points that are required for a survival plot to be produced and not seriously compromise privacy. This could be a future improvement.

Another area of future work could be generating synthetic survival data using the *dsSynthetic* package [10].

## Abbreviations

LOESS: Locally Estimated Scatterplot Smoothing.

## Declarations

### Ethics approval and consent to participate

No ethics approval and consent to participate was necessary.

### Consent for publication

Not applicable.

### Availability of data and materials

All code is available from the following repositories:

https://github.com/neelsoumya/dsSurvivalClient/

https://github.com/neelsoumya/dsSurvival/

A tutorial in bookdown format with executable code to generate plots using synthetic data is available here:

https://neelsoumya.github.io/dsSurvivalbookdown/

### Author’s contributions

SB carried out the analysis and implementation, participated in the design of the study and drafted the manuscript. TB drafted the manuscript, carried out the analysis and implementation, participated in the design of the study and directed the study. All authors read and approved the final manuscript and agree to be personally accountable for their contributions.

## Acknowledgements

We acknowledge the help and support of the DataSHIELD technical team. We are especially grateful to Demetris Avraam, Paul Burton, Stuart Wheater, Eleanor Hyde and Wolfgang Vichtbauer for fruitful discussions and feedback.

## Authors’ information

SB is a Senior Research Fellow at the University of Cambridge. He worked in industry for many years before completing a PhD in applying computational techniques to interdisciplinary topics. He has worked closely with domain experts in finance, healthcare, immunology, virology, and cell biology.

TB is a Senior Data Scientist at the University of Cambridge and has worked in industry and academia.

