## Supplementary file for "dsSurvival 2.0: Privacy enhancing survival curves for survival models in the federated DataSHIELD analysis system"

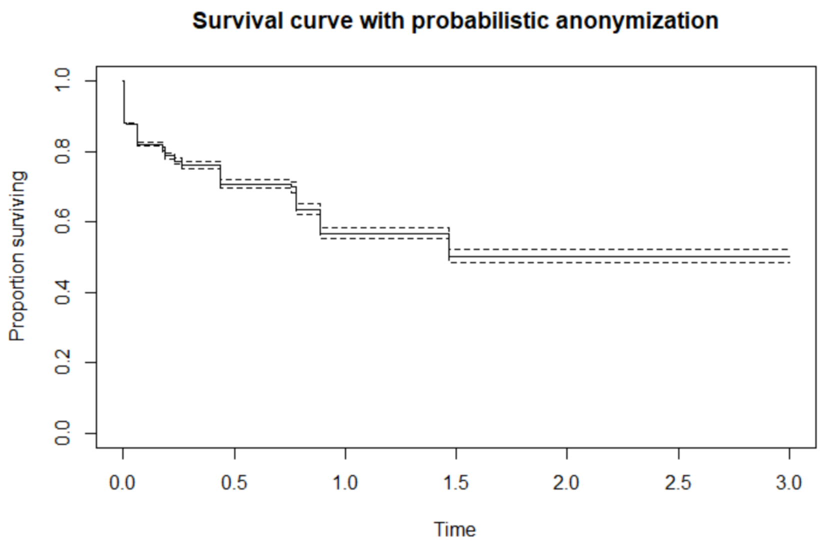


**Supplementary Figure 1.** A survival curve generated using probabilistic anonymization. This adds noise to the Y-axis (fraction survived) using probabilistic anonymization.

### Supplementary Information

In this section, we outline additional analysis that we performed.

A survival curve where we added noise to the Y-axis (fraction survived) using probabilistic anonymization is shown in Figure S1.

We conducted ablation studies to reduce the number of patients in synthetic data and observe the effect on survival curves. The survival curve for probabilistic anonymization is shown in Figure S2. This looked at a reduced number of patients (1000 as opposed to 10000 in the original dataset).

We conducted additional ablation studies where we used the automated smoothing technique on a dataset where we randomly reduced the number of patients from 137 to 25 (ablation study). The resulting privacy enhancing survival curve is shown in Figure S3.

The survival curve with one additional patient removed is shown in Figure S3.

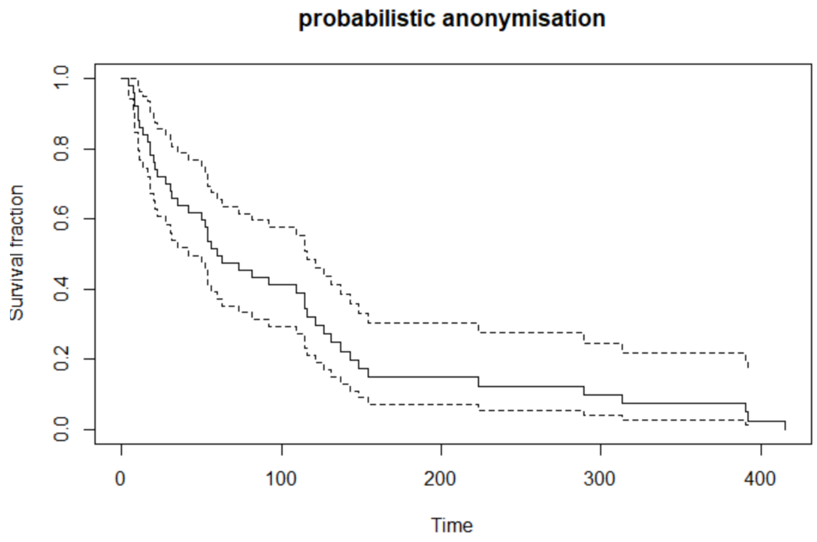


**Supplementary Figure 2.** Survival curve generated using probabilistic anonymization with fewer total number of patients (1000 as opposed to 10000 in the original data).

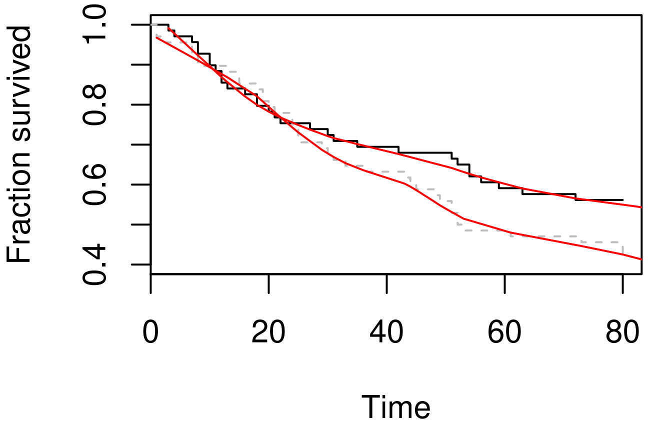


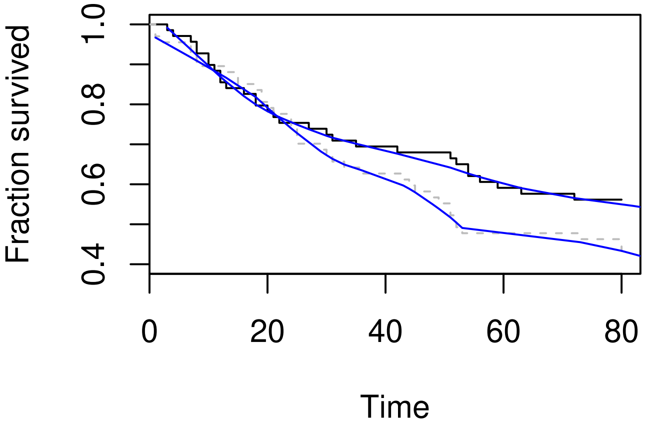


**Supplementary Figure 3.** Top panel: Survival curve with LOESS smoothing using the *fANCOVA* package with reduced number of patients (25 as opposed to 137) in the veteran dataset. The original unmodified survival curve is shown in black. Bottom panel: Survival curve with LOESS smoothing using the *fANCOVA* package with reduced number of patients (25 as opposed to 137) in the veteran dataset and one patient removed.
